# Dopaminergic Modulation of Short-Term Associative Memory in *Caenorhabditis elegans*

**DOI:** 10.1101/2025.02.20.639379

**Authors:** Anna McMillen, Caitlin Minervini, Renee Green, Michaela E. Johnson, Radwan Ansaar, Yee Lian Chew

## Abstract

Forgetting, the inability to retrieve previously encoded memories, is an active process involving neurotransmission, second messenger signalling, and cytoskeletal modifications. Forgetting is thought to be essential to remove irrelevant memories and to increase the capacity to encode new memories. Therefore, identifying key regulators of active forgetting is crucial to advance our understanding of neuroplasticity. In this study, we utilised the compact and tractable *Caenorhabditis elegans* model to investigate the role of the neurotransmitter dopamine in forgetting. We conducted butanone associative learning assays based on an established protocol and used mutant strains deficient in dopamine synthesis (tyrosine hydroxylase CAT-2 and dopamine transporter DAT-1) and signalling (G protein-coupled receptors DOP-1, DOP-2, and DOP-3) to assess the impact on learning and memory retention. Learning was measured immediately post-training, and memory retention was evaluated every 0.5 hours up to 2 hours. Our results show that animals lacking dopamine display a modest enhancement in learning relative to wild-type, with the learned association persisting for at least 2 hours after training. We also found that D2-like receptors DOP-2 and DOP-3 act together to modulate the forgetting process, with D1-like receptor DOP-1 functioning redundantly. Furthermore, re-expression of CAT-2 tyrosine hydroxylase in ADE and/or CEP neurons was unable to rescue the memory retention phenotype observed in *cat-2* mutants, suggesting that dopamine release from all dopaminergic neurons is required to modulate forgetting. These findings highlight the critical role of dopamine in forgetting, consistent with findings in *Drosophila*, and suggest potential relevance for understanding memory retention during healthy ageing and in conditions with dopamine imbalances such as Parkinson’s disease.

## Introduction

The nematode *Caenorhabditis elegans* serves as an excellent model for studying the molecular regulation of memory due to its genetic tractability, well-characterised nervous system, and capacity to learn and remember in ways analogous to higher organisms (Ardiel & Rankin 2010). In *C. elegans* and other animals, memory is categorised into short-term and long-term, primarily based on functional gene conservation with analogous processes as defined other organisms (Kandel *et al*. 2014; Brandel-Ankrapp & Arey 2023; Rahmani & Chew 2021; Giles *et al*. 2006).

On the other hand, forgetting involves the inability to retrieve previously encoded memories. Research indicates that forgetting is an active process requiring protein synthesis, and its regulation involves processes related to neurotransmission, second messenger signalling, and cytoskeletal modifications (Stein & Murphy 2014; Liu *et al*. 2022). Identifying the key modulators of active forgetting and understanding their mechanisms, distinct from those involved in learning and short- or long-term memory retention, is crucial to advance our knowledge of neuroplasticity.

Studies investigating the underlying mechanisms involved in memory regulation and forgetting have reported diverse molecular pathways. Two major regulatory mechanisms have been identified: 1) the TIR-1 and MAP kinase/JNK pathway, along with associated proteins (membrane protein MACO-1 and receptor tyrosine kinase SCD-2 and its ligand HEN-1) play roles in olfactory adaptation (Inoue *et al*. 2013; Kitazono *et al*. 2017; Arai *et al*. 2022), while 2) the RNA-binding protein MSI-1 regulates forgetting by downregulating the Arp2/3 complex (Hadziselimovic *et al*. 2014) in olfactory conditioning. Other actin regulatory proteins such as α-adducin ADD-1 and Rho family G proteins are implicated in memory loss, albeit with RAC-2 acting independently of actin dynamics (Vukojevic *et al*. 2012; Bai *et al*. 2022).

Neurotransmitters such as serotonin and acetylcholine also regulate memory retention and forgetting in *C. elegans* (Liu *et al*. 2022; Niu *et al*. 2022). In addition, dopamine plays a crucial role in modulating short-term associative memory in *C. elegans* and other organisms (McMillen & Chew 2023). Dopamine’s role in forgetting is well-documented in studies of aversive olfactory learning in *Drosophila melanogaster*: blocking dopaminergic neurons after learning enhances memory retention, while stimulation decreases or eliminates it (Berry *et al*. 2012; Berry *et al*. 2015; Berry *et al*. 2018). To determine if dopamine’s role in forgetting is conserved across species, in this study, we utilised the compact and tractable *C. elegans* to dissect the effect of dopamine on forgetting and to identify the neural circuits involved.

To do this, we conducted butanone associative learning assays according to (Kauffman *et al*. 2011) and tested mutants deficient in dopamine synthesis and signalling. Behavioural tests for forgetting in *C. elegans* involve assessing learned behaviours at various time points post-conditioning to observe when responses revert to baseline levels. Our data indicate that dopamine-deficient animals exhibit a slight improvement in learning compared to wild-type, and the memory of this learned association is retained for at least 2 hours post-training. Additionally, we found that DOP-2 and DOP-3 D2-like receptors work together to modulate the forgetting of this learned association. Lastly, CAT-2 function is required across all dopaminergic neurons to support memory retention at 2 hours post-conditioning, as re-expression in ADE alone, CEP alone, or in both ADE and CEP together was insufficient to rescue the memory defect. Taken together, our data showing that dopamine signals are required for forgetting is consistent with findings in other invertebrates (Berry *et al*. 2012). These findings highlight the critical role of dopamine in the process of forgetting and suggest implications for understanding the pathological mechanisms that contribute to cognitive deficits in neurodegenerative diseases such as Parkinson’s disease.

## Materials and Methods

### C. elegans strain maintenance

Young adult (day 1) hermaphrodite *C. elegans* were grown using standard conditions on nematode growth medium (NGM) agar in petri dishes at 22 °C for all experiments (Brenner 1974). Worms were cultured at 22 °C for at least two generations prior to all learning assays. For all experiments, animals were fed *Escherichia coli* (*E. coli*) strain OP50.

A list of strains and transgenic lines used, including full genotype information, is in **Table 1**. Genotypes of mutant strains were confirmed via PCR, based on information in Wormbase version WS295 (Sternberg *et al*. 2024). Transgenes were generated through standard microinjection procedures as in (Chew *et al*. 2018a; Chew *et al*. 2018b; Gadenne *et al*. 2022).

**Table 1:** Genotype information for strains used in this study. For transgenic lines generated in this study, numbers in square brackets refer to the concentration injected in ng/µL.

| Strain name | Genotype | Source |
| --- | --- | --- |
| Bristol N2 | Wild-type | *Caenorhabditis* Genetics Centre (CGC) |
| CB1112 | *cat-2(e1112)* | *Caenorhabditis* Genetics Centre (CGC) |
| RM2702 | *dat-1(ok157)* | *Caenorhabditis* Genetics Centre (CGC) |
| LX645 | *dop□1(vs100)* | *Caenorhabditis* Genetics Centre (CGC) |
| LX702 | *dop□2(vs105)* | *Caenorhabditis* Genetics Centre (CGC) |
| LX703 | *dop□3(vs106)* | *Caenorhabditis* Genetics Centre (CGC) |
| LX706 | *dop□1(vs100); dop□2(vs105)* | *Caenorhabditis* Genetics Centre (CGC) |
| LX704 | *dop□2(vs105); dop□3(vs106)* | *Caenorhabditis* Genetics Centre (CGC) |
| LX705 | *dop□1(vs100); dop□3(vs106)* | *Caenorhabditis* Genetics Centre (CGC) |
| LX734 | *dop□1(vs100); dop□2(vs105); dop□3(vs106)* | *Caenorhabditis* Genetics Centre (CGC) |
| YLC299 | *cat-2(e1112); bals4 [dat-1p::GFP + dat-1p::CAT-2]* | *cat-2(e1112)* crossed with UA57 from CGC |
| CLP1318 | *cat-2(n4547); twnEx593[Pace-1::cat-2; P<sub>gcy-8</sub>::mCherry]* | (Chou *et al.* 2022) |
| CLP1320 | *cat-2(n4547);twnEx599[Pexp-1::cat-2; Pgcy-8::mCherry]* | (Chou *et al.* 2022) |
| YLC381 | *dop-2(vs105)V; dop-3(vs106)X; pepEx040[Pdop-2(5kb)::dop-2S cDNA::let-858 UTR (pYLC102) [50]; Pdop-3(5kb)::dop-3a cDNA::let-858 UTR (pYLC104) [50]; ccGFP [50]]* | This study |
| YLC382 | *dop-2(vs105)V; dop-3(vs106)X; pepEx043[Pdop-2(5kb)::dop-2S cDNA::let-858 UTR (pYLC103) [50]; Pdop-3(5kb)::dop-3a cDNA::let-858 UTR (pYLC104) [50]; ccGFP [50]]* | This study |
| CLP833 | *cat-2(n4547); twnEx610[Pace-1::cat-2, Pexp-1::cat-2, Pgcy-8::mCherry]* | Gift from Chun-Liang Pan, National Taiwan University School of Medicine |

### Molecular biology

Transgenes were cloned using the NEB HiFi cloning (NEB #E2621). Promoters for the *dop-2* and *dop-3* gene were both cloned 5 kb before the ATG, starting from CAAGACGGACGCAGTCCG and CTGGAATATTCCAGAACTTTC, respectively. *dop-2* and *dop-3* cDNA were amplified from wild-type N2 RNA, extracted using the using the Monarch Total RNA Kit (NEB #T2010S) and converted to cDNA using the Luna® Universal One-Step RT-qPCR Kit (NEB #E3005S). Two forms of the *dop-2* cDNA sequence were cloned: one aligned to the DOP-2S sequence published in (Suo *et al*. 2009) (pYLC103), the other is identical to this sequence but lacks a AGTACGACGGCAAAT sequence from the 1385^th^ nucleotide position, corresponding to amino acids STTAN (pYLC102). The *dop-3* cDNA sequence aligned with the expected sequence for DOP-3, isoform a (T14E8.3a.1) (Sternberg *et al*. 2024).

### Behavioural assays

The butanone appetitive learning assay (Kauffman *et al*. 2011; Rahmani *et al*. 2024) tests naïve worms, conditioned worms that are tested directly after training (t=0), and conditioned worms that go through a ‘hold’ period between training and testing (t=0.5–2 h). Briefly, *C. elegans* populations were washed twice in M9 buffer (86 mM NaCl, 42 mM Na_2_HPO_4_, 22 mM K_3_PO_4_ (pH 6.0), 1 mM M_g_SO_4_) to constitute the ‘naïve’ group in butanone chemotaxis assays. Remaining groups were then subjected to a 1-hour starvation period in M9 buffer on a shaker (175 rpm at room temperature). Conditioned groups were then transferred onto NGM plates pre-seeded with 100 µL *E. coli* OP50 and exposed to 10% (v/v) butanone (Sigma-Aldrich, #360473; final concentration, diluted in 95% ethanol), streaked on plate lids, for 1 h. The conditioned groups undergoing the ‘hold’ period were transferred onto NGM plates pre-seeded with 100 µL *E. coli* OP50 for the assigned time periods. All groups were transferred onto salt-deficient agar in 6 cm petri dishes to perform butanone chemotaxis assays as described in Kauffman *et al*. (2010) and Rahmani *et al*. (2024). Butanone chemotaxis assay plates were prepared with two equidistant points both spotted with sodium azide, as well as 95% (v/v) ethanol on one position (control) and 10% (v/v) butanone diluted in 95% (v/v) ethanol on the other spot. Butanone chemotaxis behaviour was quantified using the following equation:

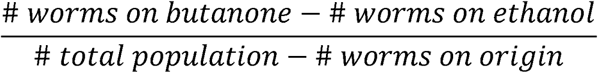

The chemotaxis index (CI) ranges between 0 (neutral butanone attraction) to 1.0 (complete butanone attraction). Each assay plate constituted one technical replicate, which each contain 50-250 worms. On very rare occasions where negative CIs are observed, these data are also excluded as these populations failed to show the standard naïve attraction to butanone. Each biological replicate constituted 3-4 technical replicates. At least three biological replicates were performed for each genotype tested, the number of biological replicates is stated for each experiment in the figure legend and are at least N>3. The Learning Index (LI) is calculated as LI _Time point_ = CI_Time point_ - CI_Naive_, where ‘Time point’ refers to the post-conditioning time point i.e. t=0, 0.5, 1, 1.5 or 2 (Kauffman *et al*. 2011).

### Statistical analysis

Statistical analysis for all experiments was performed using GraphPad Prism version 8.0. In general, where two groups were compared, a paired t-test was used. A one-way ANOVA with Dunnett’s multiple comparisons test was performed to compare post-conditioning time points with naïve chemotaxis indices (CIs) within a single genotype, or with Tukey’s multiple comparisons test to compare CIs between genotypes for naïve and t=0. The alpha value for all analyses is set at 0.05.

## Results

### Worms lacking dopamine show extended short-term associative memory

Dopamine regulates both non-associative and associative learning in *C. elegans* (McMillen & Chew 2023). To determine if dopamine regulates forgetting, we used the olfactory associative memory paradigm (**Figure 1A**) (Kauffman *et al*. 2011) to assess associative memory over time in mutants deficient in dopamine signalling. Briefly, worms are conditioned with the odour butanone (conditioned stimulus) and presence of food (unconditioned stimulus) for 1 hour on solid media, before being placed onto ‘hold’ plates containing food but no butanone for 0.5 to 2 hours. Then, worm populations are assessed for odour preference in a two-choice chemotaxis assay as in (Kauffman *et al*. 2011), where high chemotaxis indices (CIs) indicate strong butanone attraction, and CIs closer to zero represent neutral butanone attraction. ‘Naïve’ worms do not undergo conditioning and are placed onto chemotaxis plates immediately, while the ‘Timepoint 0’ (t=0) cohort are placed onto chemotaxis plates directly after conditioning. Conditioned worms undergo a 1 hour ‘pre-starvation’ step to increase the value of the unconditioned stimulus (food) (Kauffman *et al*. 2011) (**Figure 1Aii)**. To simplify the visualisation of behavioural assay data, we only display comparisons between genotypes at each post-conditioning time point, and between naive and t=0 for wild-type and other genotypes to show if learning has occurred. Other statistical comparisons, specifically those that compare chemotaxis behaviour WITHIN genotypes (i.e. between naïve and post-conditioning time points) are described in the text. After conditioning (t=0), wild-type worms show an increased attraction to butanone (shown by a higher chemotaxis index, CI) compared with the naïve cohort, indicating the capacity for associative olfactory learning. This memory decreases over time, returning to baseline (naïve) levels after ∼0.5 hours (comparison between CIs of naïve and 0.5 h post-conditioning cohorts: p-value = 0.6754; one-way ANOVA with Dunnett’s post-test) (**Figure 1B**).

**Figure 1:**
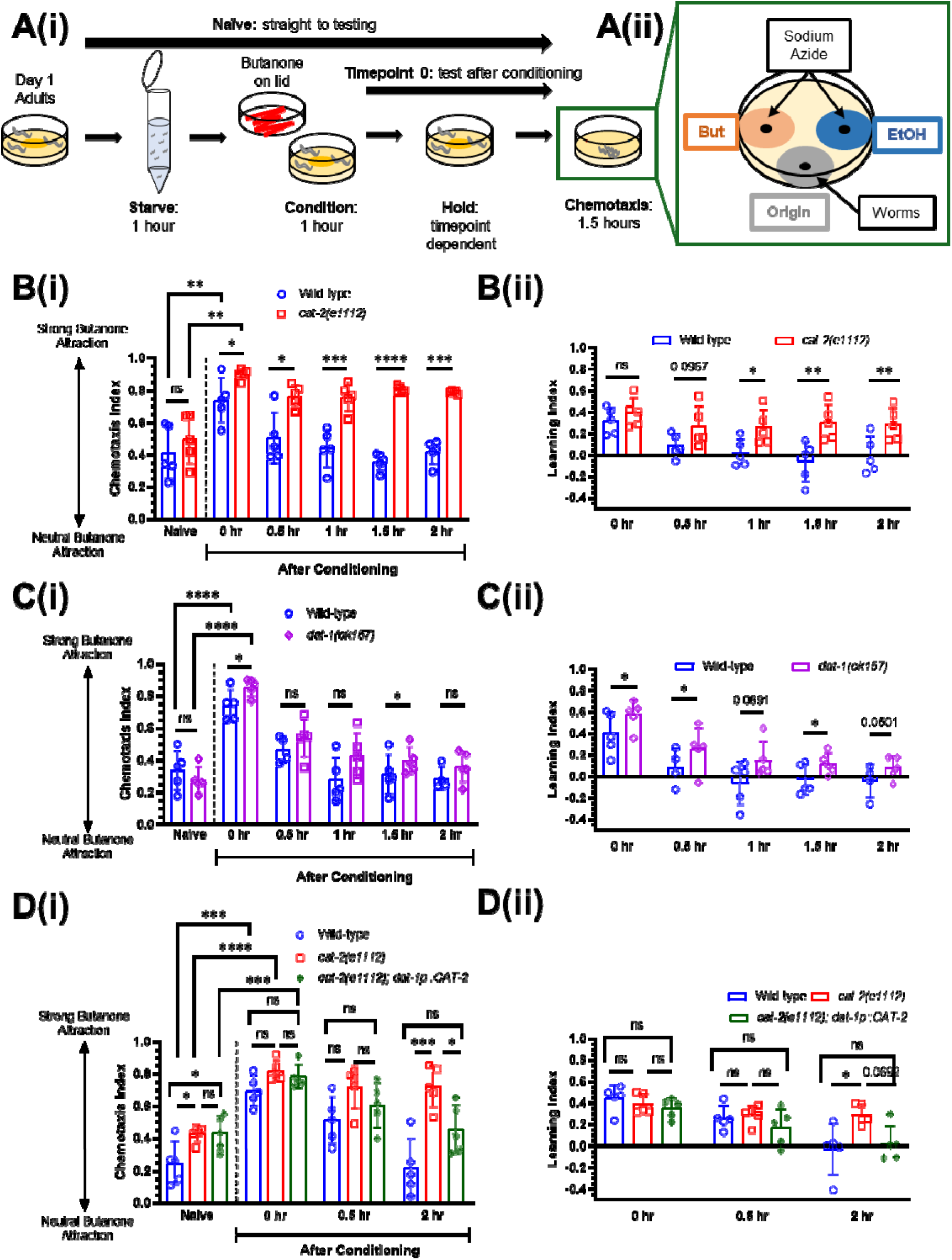
Worms lacking dopamine show extended short-term associative memory. **A) (i)** Schematic of butanone associative learning and short-term memory assay, adapted from (Kauffman *et al*. 2011). Naïve worms are placed directly onto chemotaxis assay plates, while conditioned worms are first starved for 1 hour in M9 buffer prior to conditioning on plates with food and butanone (streaked on lid). After conditioning, worms are transferred to ‘hold’ plates with food (no odour) for 0.5–2 hours before being placed onto chemotaxis plates. **(ii)** Layout of chemotaxis assay plate indicating where butanone (But), ethanol (EtOH) and sodium azide are spotted. Worms are placed on the ‘origin’, as indicated. **B)** Wild-type and *cat-2(e1112)* mutant worms tested for butanone associative memory (*n =* 5 biological replicates) **(i)** chemotaxis index and **(ii)** learning index. **C) (i)** Wild-type and *dat-1(ok157)* mutant worms tested for butanone associative memory (*n =* 5 biological replicates) **(i)** chemotaxis index and **(ii)** Learning index. **D) (i)** Wild-type and *cat-2* ‘rescue’ worms re-expressing CAT-2 in the dopaminergic neurons tested for butanone associative memory (*n =* 4 biological replicates). **(ii)** Learning indices for wild-type, *cat-2(e1112)* mutant worms, and CAT-2 rescue worms, calculated from **D(i)**. For butanone assays (**B, C, D**), graphs show 4-5 biological replicates: each data point represents one biological replicate (which includes four technical replicates). Each technical replicate consists of 50-250 worms. Behaviour was quantified using chemotaxis indices (CI) for naïve/untrained worms and conditioned animals at 0, 0.5-, 1-, 1.5-and 2-hours post-conditioning. Statistical analysis: a pairwise t-test was performed to compare genotypes for each time point for **B** and **C**, and a one-way ANOVA with Tukey’s multiple comparisons test was performed to compare post-conditioning t=0 with naïve chemotaxis indices (CIs) between genotypes for all graphs and to compare genotypes for each time point for **D**. Error bars = mean ± SD. p-values are represented by **** ≤ 0.0001; *** ≤ 0.001, ** ≤ 0.01; * ≤ 0.05; ns = non-significant.

We first tested the *cat-2* tyrosine hydroxylase mutant strain that lacks the enzyme required to synthesise dopamine (Lints & Emmons 1999). These worms have substantially reduced dopamine levels (Sanyal *et al*. 2004). Interestingly, *cat-2* mutant worms show a small but significant improvement in learning, and an increased attraction to butanone for up to 2 hours (within genotype comparison between CIs of *cat-2* naïve and 2 h post-conditioning cohorts: p-value <0.0001; one-way ANOVA with Dunnett’s post-test), suggesting they retain the memory of the learnt association for a longer period compared with wild-type controls (**Figure 1B**). Thus, for the purposes of our study, we consider forgetting to be the loss of the learnt phenotype (increased attraction to butanone) at 2 hours post-conditioning. We also calculated the learning index (LI), defined as the CI at the post-conditioning time point minus the CI of the naïve cohort, for all time points tested. This is shown for data from wild-type and *cat-2* mutant worms in **Figure 1Bii**. The *cat-2* mutant strain shows a higher LI for t=1, 1.5 and 2 compared with wild-type controls, consistent with a longer retention of the learnt phenotype at later time points post-conditioning.

To determine if a milder reduction in dopamine has the same effect, we next tested *dat-1* mutant worms with a mutation in the DAT-1 sodium-dependent dopamine transporter. These worms are able to synthesise dopamine; however, dopamine reuptake from the synaptic cleft is impaired, reducing the amount of residual dopamine in the nervous system (Formisano *et al*. 2020; Kindt *et al*. 2007). *dat-1* mutant worms showed improved learning compared with wild-type controls immediately after conditioning (t=0) and retained memory of the learnt behaviour at t=0.5 (within-genotype comparison between naïve and t=0.5 cohorts: p-value <0.01 for *dat-1*; one-way ANOVA with Dunnett’s post-test). Comparing CIs indicates that *dat-1* mutant worms do not display an extended memory phenotype like *cat-2* mutants (within-genotype comparisons between naïve and time points after t=0.5: all p-values >0.05; one-way ANOVA with Dunnett’s post-test) (**Figure 1Ci**). Notably, the *dat-1* mutant strain shows a significantly higher LI at t=1.5, and close to significantly different at t=1 and t=2 compared with wild-type controls (**Figure 1Cii**), suggesting that there is an effect on memory, albeit less pronounced than *cat-2* mutants. This indicates that while excess dopamine in the synaptic cleft (as seen in *dat-1* mutants) leads to a memory retention phenotype at 1.5 hours, dopamine synthesised by CAT-2 is required for normal retention of the learnt phenotype (i.e. the loss of memory) at 2 hours post-conditioning.

Finally, we tested if re-expression of CAT-2 in dopaminergic neurons (using the *dat-1* promoter) rescues the extended memory phenotype in *cat-2* mutants. We found that transgenic worms re-expressing CAT-2 generally phenocopied wild-type animals: the re-expression strain rescued the *cat-2* phenotype to wild-type levels at t=2 post-conditioning, as shown by comparing CIs to the *cat-2* mutant (**Figure 1Ci**). Comparing LIs between the rescue strain and the mutant strain was near-significant at t=2, with p-value 0.069 (**Figure 1Cii**). This indicates that the enzyme required for dopamine synthesis, CAT-2, is required for forgetting and acts in the dopaminergic neurons.

### Dopamine receptors act redundantly to modulate short-term memory

In *C. elegans*, dopamine receptors are categorised as ligand-gated ion channels (LGIC) or G protein-coupled receptors (GPCR). Dopaminergic GPCRs are divided into D1-like and D2-like receptors, which stimulate or inhibit adenylyl cyclase, respectively (Chase & Koelle 2007; Girault & Greengard 2004; Mersha *et al*. 2013). DOP-1 is a D1-like receptor, while DOP-2 and DOP-3 are D2-like receptors (Chase & Koelle 2007; Mersha *et al*. 2013) (**Figure 2A**). We first tested *dop-1, dop-2* and *dop-3* single mutants to determine if they showed a memory retention phenotype similar to *cat-2* mutant worms. We found that none of the single mutants showed statistically significant learning or memory differences compared with wild-type controls (**Figure 2B, Figure S1**): CIs and LIs for all mutants showed intact learning at t=0, and butanone preference had returned to baseline (naïve) levels at t=0.5 (within genotype comparisons for all dopamine receptor mutants between naïve and t=0.5: p-values >0.05 for all genotypes; one-way ANOVA with Dunnett’s post-test). This indicates that dopamine does not regulate forgetting by acting through DOP-1, DOP-2 or DOP-3 alone.

**Figure 2:**
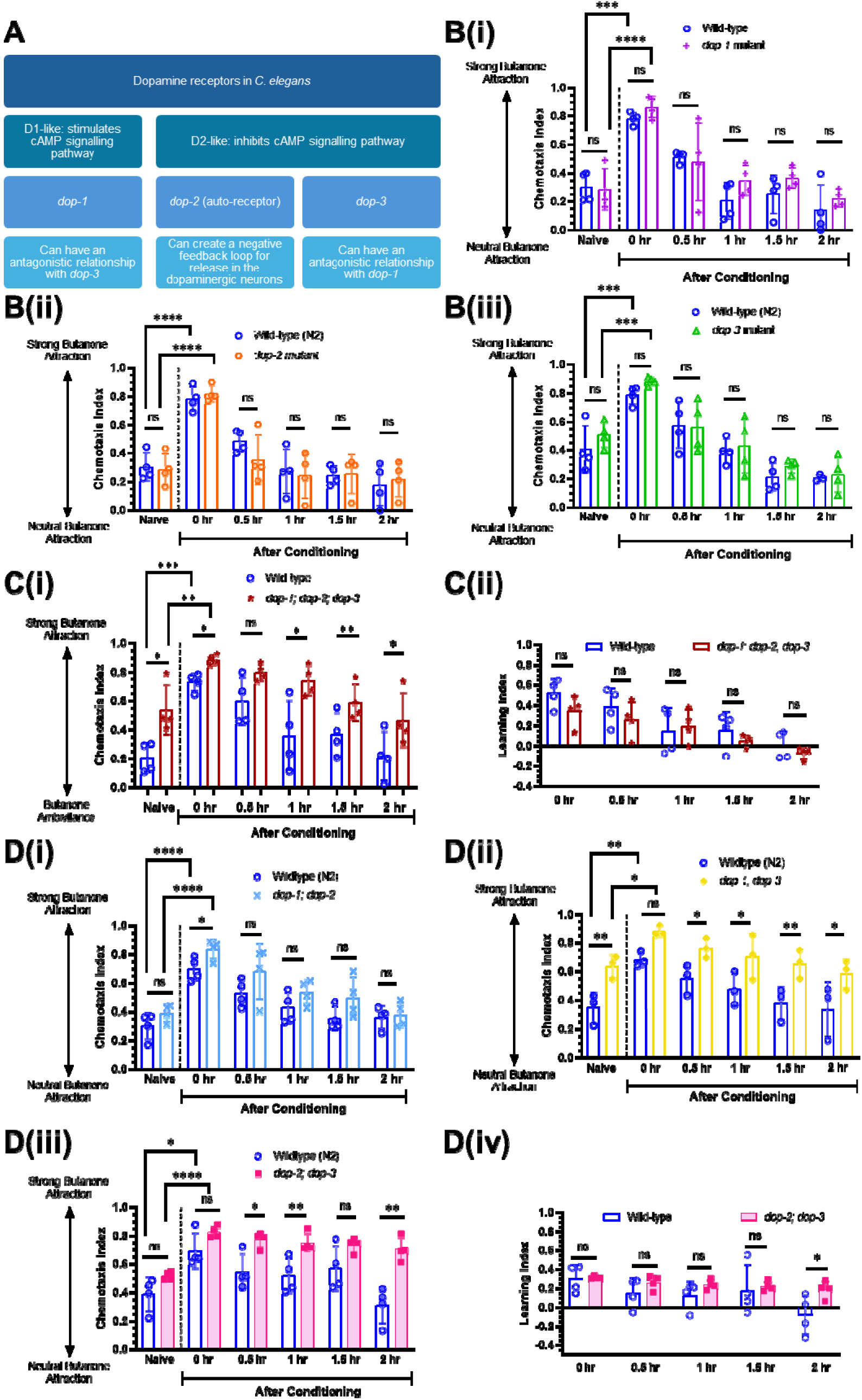
Dopamine receptors act synergistically to modulate short-term memory. **A)** *C. elegans* dopamine receptors DOP-1, DOP-2, and DOP-3 are D1 or D2-like receptors (Chase & Koelle 2007; Mersha *et al*. 2013). **B)** Butanone associative memory assay for wild-type worms and worms containing single mutations in **(i)** *dop*lJ*1(vs100)* (*n =* 4 biological replicates), **(ii)** *dop*lJ*2(vs105)* (*n =* 4 biological replicates), and **(iii)** *dop*lJ*3(vs106)* (*n =* 4 biological replicates). **C)** Butanone associative memory assay for wild-type and *dop*lJ*1;dop*lJ*2;dop*lJ*3* triple mutant worms (*n =* 4 biological replicates) **(i)** chemotaxis index and **(ii)** learning index **D)** Butanone associative memory assay for wild-type and **(i)** *dop*lJ*1;dop-2* (*n =* 4 biological replicates), **(ii)** *dop*lJ*1;dop-3* (*n =* 3 biological replicates), and **(iii)** *dop*lJ*2;dop-3* (*n =* 4 biological replicates) double mutant worms. **(iv)** The learning index calculated from **(iii)**. For butanone assays (**B, C, D**), each data point represents one biological replicate (which includes four technical replicates). Each technical replicate consists of 50-250 worms. Behaviour was quantified using chemotaxis indices (CI) for naïve/untrained worms and conditioned animals at 0, 0.5-, 1-, 1.5- and 2-hours post-conditioning. Statistical analysis: a pairwise t-test was performed to compare genotypes for each time point, and a one-way ANOVA with Tukey’s multiple comparisons test was performed to compare post-conditioning t=0 with naïve chemotaxis indices (CIs) between genotypes. Error bars = mean ± SD. p-values are represented by **** ≤ 0.0001; *** ≤ 0.001, ** ≤ 0.01; * ≤ 0.05; ns = non-significant.

One possibility is that these receptors could be acting in a functionally redundant manner for dopamine signalling to modulate forgetting in *C. elegans*, as in other behaviours such as bacterial avoidance (Chou *et al*. 2022) or the escape response to olfactory cues (Baidya *et al*. 2014). Therefore, we next tested a *dop-1;dop-2;dop-3* triple mutant strain and found that they displayed similar memory compared with wild-type controls. Comparisons between naïve and post-conditioning CIs and LIs for all time points within the triple mutant strain were not statistically significant (for CIs, all within genotype comparisons are p>0.05, one-way ANOVA with Dunnett’s post-test) (**Figure 2Ci, 2Cii**). We postulated that there could be a synergistic or redundant effect between D1 and D2-like dopamine receptors that could be masking the effects of individual receptors (Chase *et al*. 2004), and so we subsequently tested double mutants for *dop-1;dop-2*, *dop-1;dop-3*, and *dop-2;dop-3* for their capacity to retain memory over time (**Figure 2D**). As in other assays, we found that wild-type controls returned to baseline (naïve) CI values by t=0.5 (or t=1 for **Figure 2Di**). This was also the case for *dop-1;dop-2* (**Figure 2Di**) and *dop-1;dop-3* (**Figure 2Dii**) double mutants, which returned to baseline CI levels by t=1 and t=0.5, respectively. In contrast, the *dop-2;dop-3* double mutants showed elevated CIs post-conditioning that had not returned to baseline by t=2 (within genotype comparisons between naïve and t>0: p-values <0.01 for all time points; one-way ANOVA with Dunnett’s post-test) (**Figure 2Diii**). Consistent with this, the LI for the *dop-2;dop-3* double mutant is significantly higher than wild-type controls at t=2 (**Figure 2Div**). Taken together, these data suggest that the D1-like receptor DOP-1 and D2-like receptors DOP-2 and DOP-3 act redundantly to modulate forgetting of an olfactory associative memory.

### Dopamine acts in dop-2/dop-3 expressing neurons to modulate short-term memory

Our data suggests that dopamine acts through D2-like dopamine GPCRs, DOP-2 and DOP-3, to promote forgetting in *C. elegans*. We re-expressed DOP-2 and DOP-3 in *dop-2;dop-3* double mutants using a transgene that contains DOP-2 cDNA (corresponding to DOP-2S in (Suo *et al*. 2009)) and DOP-3a cDNA, expressed under the control of 5 kb of promoter sequence for both *dop-2* and *dop-3* (see Materials and Methods for details). Due to practical limitations on the number of groups that can be handled in a single experiment, for these assays we tested only naïve and post-conditioning t=0, 0.5 and 2 h cohorts for four genotypes: wild-type, *dop-2;dop-3* double mutants, and two versions of the transgenic DOP-2/3 rescue line (**Figure 3A** and **B**). As the DOP-2/3 rescue transgenic lines contain extrachromosomal arrays, there may be minor differences in transgene expression, so we tested two versions to assess the role of DOP-2 and DOP-3 in forgetting. We found for data shown in both **Figure 3A** and **3B** that, as expected, wild-type controls showed intact learning at t=0, as indicated by higher CI compared with naïve worms, and that this had returned to baseline at t=2 (comparisons between naïve and t=0, 0.5, and 2 for all genotypes are indicated underneath each graph). Consistent with **Figure 2Diii**, *dop-2;dop-3* mutants also displayed intact learning at t=0, and the memory of this learnt association did not return to baseline levels at t=2 (comparison between CIs for naïve and t=2: p-value <0.05; one-way ANOVA with Dunnett’s post-test) (**Figures 3A, 3B**), indicating an extended memory retention phenotype. When comparing CIs, we found that both DOP-2/3 re-expression lines showed a trend towards a rescue phenotype: transgenic worms showed intact learning at t=0, and had returned to naïve levels at t=2 (within-genotype comparison between CIs for naïve and t=2: p-value >0.05; one-way ANOVA with Dunnett’s post-test) (**Figures 3Ai, 3Bi**). Although the mean CI for both rescue strains was intermediate between wild-type or *dop-2;dop-3* mutant controls, the CIs of DOP-2/3 rescue worms were not significantly different from either of the controls at t=2 (**Figures 3Ai, 3Bi**). Comparison of LIs at t=2 showed that the LIs of both rescue strains were lower than the mutant (approaching wild-type levels), but this was not statistically significant (**Figures 3Aii, 3Bii**); notably, the comparison between the rescue strain and mutant was near-significant for v2 at p=0.07 (**Figure 3Bii**). Taken together, these data suggest a consistent trend suggesting that re-expression of DOP-2 and DOP-3 in the *dop-2;dop-3* double mutant background contributes towards restoring the forgetting phenotype. While the rescue effect does not reach statistical significance, the observed effects support a potential role for these receptors in modulating memory decay.

**Figure 3:**
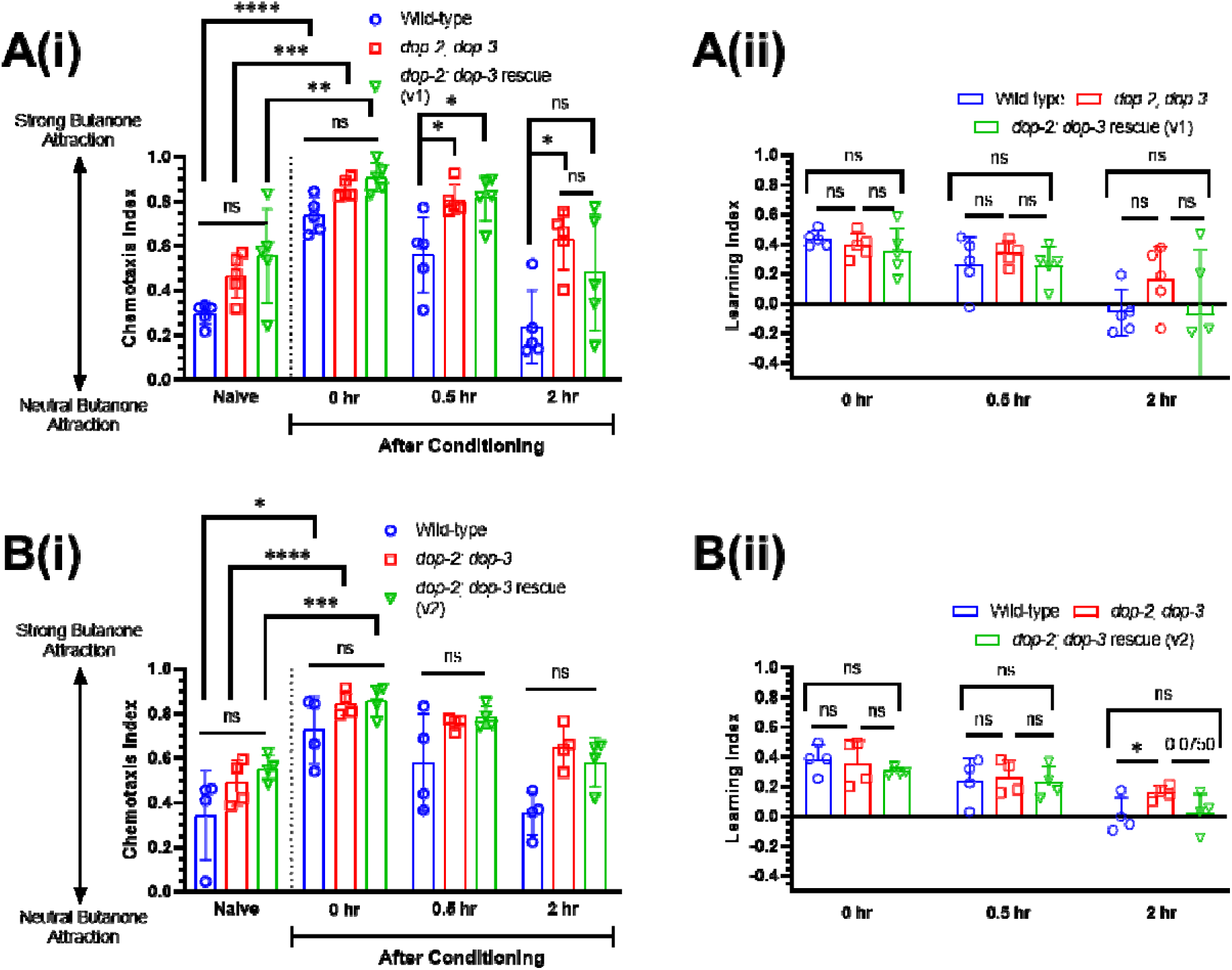
Dopamine acts through DOP-2 and DOP-3 to modulate short-term memory. **A)** Butanone associative memory assay for wild-type worms, *dop-2;dop-3* double mutant worms and *dop-2;dop-3* double mutant worms re-expressing DOP-2 and DOP-3 under the control of their endogenous promoters, **(i)** the chemotaxis index and **(ii)** the learning index graph. DOP-2 in this transgene is a truncated version missing 5 amino acids when compared to the DOP-2S sequence (*n =* 5 biological replicates). **B)** Butanone associative memory assay for wild-type worms, *dop-2;dop-3* double mutant worms and *dop-2;dop-3* double worms re-expressing DOP-2 and DOP-3 under the control of their endogenous promoters; **(i)** the chemotaxis index and **(ii)** the learning index graph. DOP-2 in this transgene exactly matches the published DOP-2S sequence (Suo *et al*. 2009) (*n =* 4 biological replicates). For rescue lines expressin extrachromosomal arrays, only CIs from transgenic worms are displayed here. Each data point represents one biological replicate (which includes four technical replicates). Each technical replicate consists of 50-250 worms. Behaviour was quantified using chemotaxis indices (CI) for naïve/untrained worms and conditioned animals at 0, 0.5-, and 2-hours post-conditioning. Statistical analysis: one-way ANOVA with Dunnett’s multiple comparisons test was performed to compare post-conditioning time points with naïv chemotaxis indices (CIs) within a single genotype (shown in tables below each graph) and with Tukey’ multiple comparisons test to compare CIs for naïve vs t=0 between genotypes. Error bars = mean ± SD. p-values are represented by **** ≤ 0.0001; *** ≤ 0.001, ** ≤ 0.01; * ≤ 0.05; ns = non-significant.

### Dopamine release through ADE and CEP neurons promotes forgetting

There are eight dopaminergic neurons in hermaphrodite *C. elegans*: two ADE neurons, four CEP neurons, and two PDE neurons (**Figure 4A**), each of which have specific synaptic and neuromodulatory inputs/outputs to other parts of the nervous system. To investigate whether specific connections are required for memory retention, we examined the role of individual subsets of dopamine-releasing neurons in the process of forgetting. To do this, we assayed transgenic lines in which CAT-2 is specifically re-expressed in the ADE (using an *exp-1* promoter) or CEP neurons (using an *ace-1* promoter), or both, in a *cat-2* mutant background (Chou *et al*. 2022). We first examined the role of ADE neurons in forgetting (**Figure 4B**): wild-type controls displayed the expected phenotype of showing an increased CI (indicating intact learning) at t=0, which had returned to baseline at t=0.5 (within genotype comparison between CIs for naïve and t=0.5: p-value >0.05; one-way ANOVA with Tukey’s post-test). We found that ADE::CAT-2 rescue worms showed intact learning at t=0, which had also returned to baseline at t=0.5 (within genotype comparison between CIs for naïve and t=0.5: p-value >0.05; one-way ANOVA with Tukey’s post-test) (**Figure 4A**). To assess whether re-expression in ADE neurons alone rescued the forgetting phenotype in *cat-2* mutants, we compared CIs and LIs at t=2 between the ADE rescue strain and mutant siblings. We found that there was no statistically significant difference in CI (**Figure 4Ai**) or LI (**Figure 4Aii**), suggesting that CAT-2 expression in ADE neurons is insufficient for normal forgetting.

**Figure 4:**
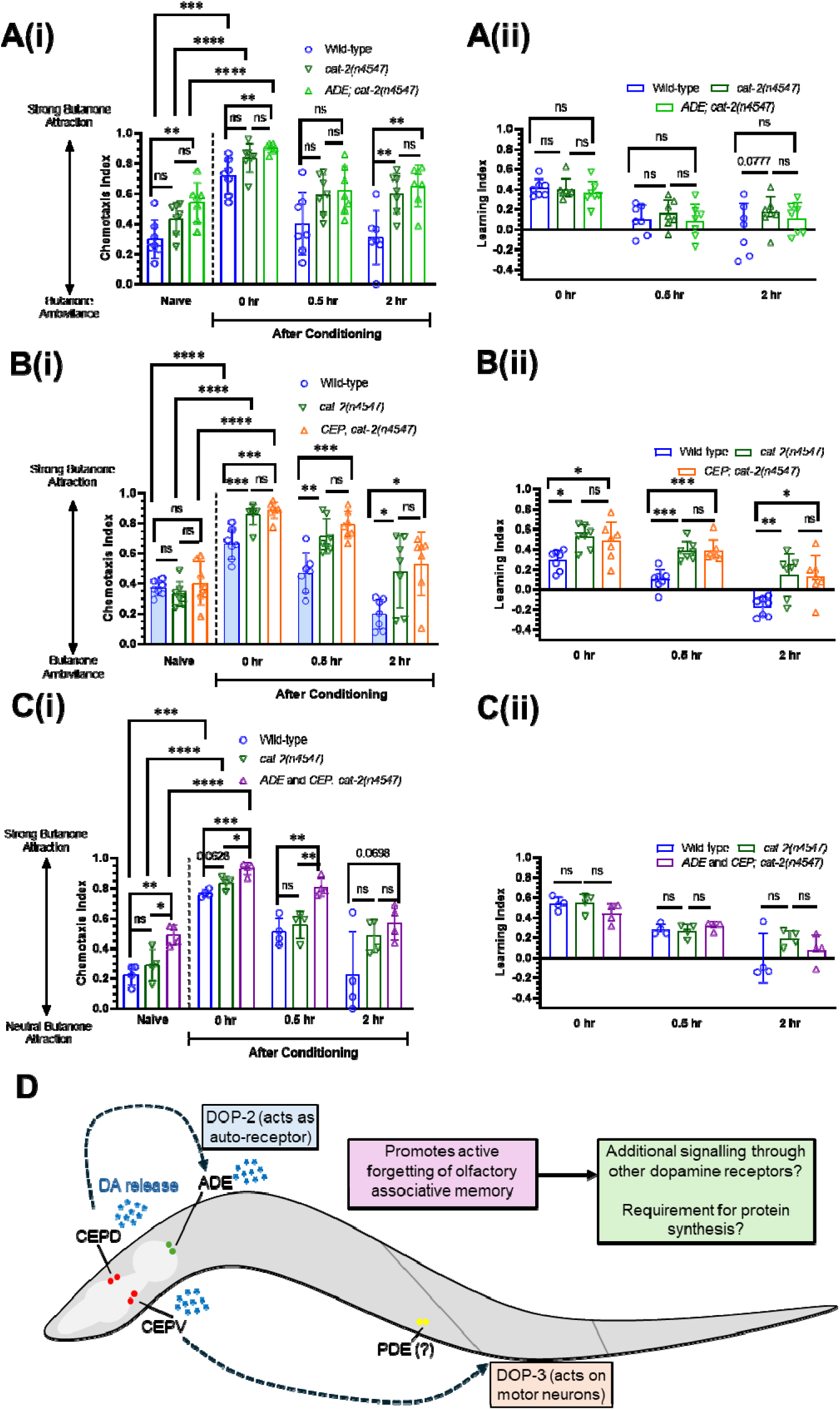
Dopamine release from ADE and CEP neurons promotes forgetting. **A)** Butanone associative memory assay for wild-type worms and *cat-2* mutant worms re-expressing CAT-2 in the ADE neurons (*n =* 7 biological replicates); **(i)** the chemotaxis index and **(ii)** the learning index graphs. **B)** Butanone associative memory assay for wild-type worms and *cat-2* mutant worms re-expressing CAT-2 in the CEP neurons (*n =* 7 biological replicates); **(i)** the chemotaxis index and **(ii)** the learning index graphs. **C)** Butanone associative memory assay for wild-type worms and *cat-2* mutant worms re-expressing CAT-2 in both the ADE and CEP neurons (*n* = 4 biological replicates); **(i)** the chemotaxis index and **(ii)** the learning index graphs. Each data point represents one biological replicate (which includes four technical replicates). Each technical replicate consists of 50-250 worms. Behaviour was quantified using chemotaxis indices (CI) for naïve/untrained worms and conditioned animals at 0, 0.5-, and 2-hours post-conditioning. Statistical analysis: a pairwise t-test was performed to compare genotypes for each time point, and a one-way ANOVA with Tukey’s multiple comparisons test was performed to compare post-conditioning t=0 with naïve chemotaxis indices (CIs) between genotypes. Error bars = mean ± SD. p-values are represented by **** ≤ 0.0001; *** ≤ 0.001, ** ≤ 0.01; * ≤ 0.05; ns = non-significant. **D)** Schematic of potential mechanism, summarising data in Figures 1-4: our findings suggest that newly synthesised dopamine from all dopaminergic neurons is required for forgetting of olfactory associative memory, acting primarily through DOP-2 and DOP-3 receptors.

We next tested CEP::CAT-2 rescue worms (**Figure 4B**) in the butanone associative memory assay. We found that wild-type worms showed the expected phenotype of intact learning at t=0 and return to baseline (naïve) CI at t=0.5. In contrast, CEP::CAT-2 worms showed significantly higher CIs at t=0 (indicating intact learning) and t=0.5 (within genotype comparison between naïve and t=0.5 gives p-value of 0.0103; one way ANOVA, Tukey’s post-test). Similarly, to assess whether re-expression in CEP neurons alone rescued the forgetting phenotype in *cat-2* mutants, we compared CIs and LIs at t=2 between the rescue strain and mutant siblings. There was no statistically significant difference in CI (**Figure 4Bi**) or LI (**Figure 4Bii**) t=2, indicating that CAT-2 expression in CEP neurons is insufficient for normal forgetting.

As re-expression of CAT-2 in ADE or CEP did not rescue the extended memory phenotype observed in *cat-2* mutants, we tested a strain where CAT-2 was re-expressed in both ADE and CEP neurons (referred to as ADE+CEP; **Figure 4C**). Similar to ADE and CEP alone, this ADE+CEP rescue line did not show a statistically different CI (**Figure 4Ci**) or LI (**Figure 4Cii**) compared to the *cat-2* mutant siblings at t=2. Taken together, this indicates that CAT-2 expression in ADEs and CEPs does not rescue the memory phenotype, and suggests that dopamine release from all dopaminergic neurons (ADEs, CEPs and PDEs), or from PDE alone, is required for normal forgetting.

## Discussion

Our study demonstrates that dopamine is essential for active forgetting in *C. elegans*. Our findings indicate that dopamine-deficient mutants, particularly *cat-2*, exhibit enhanced memory retention following conditioning, consistent with a delayed forgetting phenotype. The *dat-1* mutant, which retains dopamine synthesis but has impaired reuptake, shows a similar trend, albeit less pronounced. These results suggest that even partial restoration of dopamine may influence forgetting, consistent with the idea that boosting dopamine levels (such as via pharmacological pretreatment) could restore the delayed forgetting phenotype in *cat-2* mutants. Additionally, our findings indicate that D2-like receptors, specifically DOP-2 and DOP-3, are required for forgetting. Finally, we observed that CAT-2 function is required across all dopaminergic neurons to support memory retention at 2 hours post-conditioning. Re-expression of *cat-2* in ADE alone, CEP alone, or in both ADE and CEP together was insufficient to rescue the memory defect observed in the mutant background at this time point (**Figure 4**).

The role of dopamine in learning and memory is conserved across species. However, while dopamine synthesis and signalling pathways are highly conserved, dopaminergic circuits between *C. elegans* and other organisms differ. In *C. elegans*, eight dopaminergic neurons act ‘extra-synaptically’ on over 100 targets, mainly motor neurons (Bentley *et al*. 2016). In mammals, dopaminergic neurons are concentrated in the ventral midbrain (SNc and VTA) and project to the basal ganglia, amygdala, and prefrontal cortex. In *Drosophila melanogaster*, small populations of dopaminergic neurons are found throughout the fly brain, with projections innervating almost the entire brain, with dense projections to the mushroom body lobes, which regulate olfactory learning (Riemensperger *et al*. 2005; Marquis & Wilson 2022). Despite these differences, our finding that dopamine is required for forgetting aligns with previous research on the role of dopamine in forgetting in *Drosophila*. In flies, blocking synaptic output from dopaminergic neurons promotes memory retention (i.e. reduces forgetting), whereas activating dopamine neurons decreases memory (Berry *et al*. 2012). Indeed, dopamine is essential for both forgetting and memory acquisition in flies, but this involves different receptors: DAMB for forgetting and dDA1 for acquisition (Berry *et al*. 2012; Himmelreich *et al*. 2017). In our study, we observed a small but robust learning improvement in *cat-2* and *dat-1* mutant worms, as well as in *dop-1;dop-2;dop-3* triple mutants (double mutant combinations of these three receptors also showed a learning improvement, albeit slightly above the alpha value of p = 0.05). This indicates that the requirement for dopamine in learning and forgetting is conserved between flies and worms, suggesting it is evolutionary important and likely also retained in higher organisms.

Our findings contrast with the work by Raj and Thekkuveettil (Raj & Thekkuveettil 2022), which showed that dopamine deficient *cat-2* mutant *C. elegans* display both a learning defect and a memory defect, i.e. they forget more quickly than wild-type controls. Although both our study and the 2022 study use the butanone associative learning assay, worms in this research were conditioned with agar plates seeded 1-2 days in advance with 100 µL OP50 (and thus already dried on the agar), whereas the Raj and Thekkuveettil study involved conditioning in 400 µL of OP50 pipetted onto unseeded agar plates immediately after worms are added. This means that the worms are being conditioned in liquid, which may influence other locomotion pathways in which dopamine plays an important role, such as in regulating “swimming” in liquid, a distinct form of locomotion separate to crawling on solid media. Dopamine modulates swimming behaviour (Xu *et al*. 2021), swimming-induced paralysis (McDonald *et al*. 2007), and swim-to-crawl transitions when moving from liquid to solid media (Vidal-Gadea *et al*. 2011). As learning is a multifaceted process modulated by multiple internal and external cues, it is not surprising that dopamine could impact memory in different ways depending on the mode of conditioning, for example due to an interaction between the response to the food and odour cues, and the swimming activity. This underscores dopamine’s multifaceted role in modulating behaviour and memory in *C. elegans*.

Our data show that both DOP-2 and DOP-3 D2-like receptors are required to modulate forgetting. Single mutants of either receptor show memory retention capacity similar to wild-type, while the *dop-2;dop-3* double mutant show enhanced memory retention similar to *cat-2* mutant animals. Taken together, our data suggest that the D1-like receptor DOP-1 and D2-like receptors DOP-2 and DOP-3 act redundantly to modulate forgetting of an olfactory associative memory. Although D1- and D2-like receptors are known to antagonise each other in signal transduction, and their opposing effects in *C. elegans* have been previously reported in dietary restriction-mediated lifespan extension (Jiang *et al*. 2022), swimming (Xu *et al*. 2021), and basal slowing behaviour in response to a food lawn (Chase *et al*. 2004), this did not appear to be the case in our study. We did not test the invertebrate specific DOP-4 receptor or dopamine ligand-gated ion channels such as LGC-53; therefore, a role for these receptors in dopamine-dependent short-term associative memory in *C. elegans* remains unknown. Importantly, although re-expression of DOP-2 and DOP-3 in the mutant background showed trends toward restoring normal forgetting, these effects did not reach statistical significance (**Figure 3**). Future studies could refine the re-expression strategy, such as by modulating DNA dosage for microinjection or selecting alternative isoforms, to enhance rescue efficacy and more clearly define the contributions of these receptors to forgetting.

Short- and long-term memory differ by the mechanism for encoding and time taken to fully decay. In *C. elegans*, short-term memory is typically considered to be relevant at 1-2 hours post-learning, while long-term memory is evaluated 12-24 hours post-learning (Rahmani & Chew 2021; Kauffman *et al*. 2011). In our study, we tested short-term associative memory only up to 2 hours post-conditioning. At this time point, dopamine-deficient *cat-2* mutants, as well as *dop-2;dop-3* double mutants, still showed robust memory. Identifying when the memory decays to baseline levels may provide further understanding of the role of dopamine in forgetting, such as whether processes associated with long-term memory e.g. *de novo* protein synthesis (Fioriti *et al*. 2015), are required. The mechanistic details of this process warrant further investigation in future studies.

In conclusion, our study leverages the worm’s compact nervous system and genetic tractability to provide new insights into the role of dopamine in forgetting. Other studies on olfactory paradigms in worms have identified diverse regulators of forgetting, indicating that multiple pathways and mechanisms are involved in this complex process. Our study adds to this body of knowledge by highlighting the specific role of dopamine and its receptors. In addition, these findings are consistent with research in *Drosophila*, suggesting a mechanism for forgetting that is conserved in even more complex brains. Based on this study, future research could focus on exploring the conserved mechanisms of forgetting across species and investigating potential therapeutic targets for memory-related disorders.

## Acknowledgements

Many thanks go to our team members in the Worm Neuroscience lab (Flinders) for thoughtful discussions, and to our colleagues at Flinders University – Dr Nicholas Eyre, Professor Damien Keating, and Professor Kim Hemsley – for feedback and scientific advice. We especially thank Pasquale Vitaro (Summer Scholar 2025) for technical assistance with behavioural assays, and Aelon Rahmani for assistance with genotyping. We are grateful to Prof Chun-Liang Pan (National Taiwan University School of Medicine) for strains CLP1318, CLP1320, and CLP833 (cell-specific rescue of *cat-2*), and Prof Satoshi Suo (Saitama Medical University, Japan) for plasmids encoding *dop-2* cDNA. Finally, we gratefully acknowledge the *Caenorhabditis* Genetics Centre, which is supported by the National Institutes of Health (P40 OD010440), for providing some of the strains used in this study (see **Table 1**).

## Author contributions

Conceptualisation/Methodology– **AM, YLC**. Data curation/ Formal analysis/ Investigation/ Resources – **AM, CM, MEJ, RG, RA, YLC.** Project administration/ Supervision/ Funding acquisition – **YLC.** Visualisation/ Writing – original draft - **AM, YLC** Writing – review & editing – **AM, MEJ, YLC**

## Data Availability Section

This study includes no data deposited in external repositories. Reagents including *C. elegans* strains and expression plasmids are available from the corresponding author upon request.

## List of abbreviations

*C. elegans*: *Caenorhabditis elegans*
CI: Chemotaxis Index
*E. Coli*: *Escherichia coli*
LI: Learning index
PCR: Polymerase Chain Reaction

## Supplementary Figure legends

**Figure S1:**
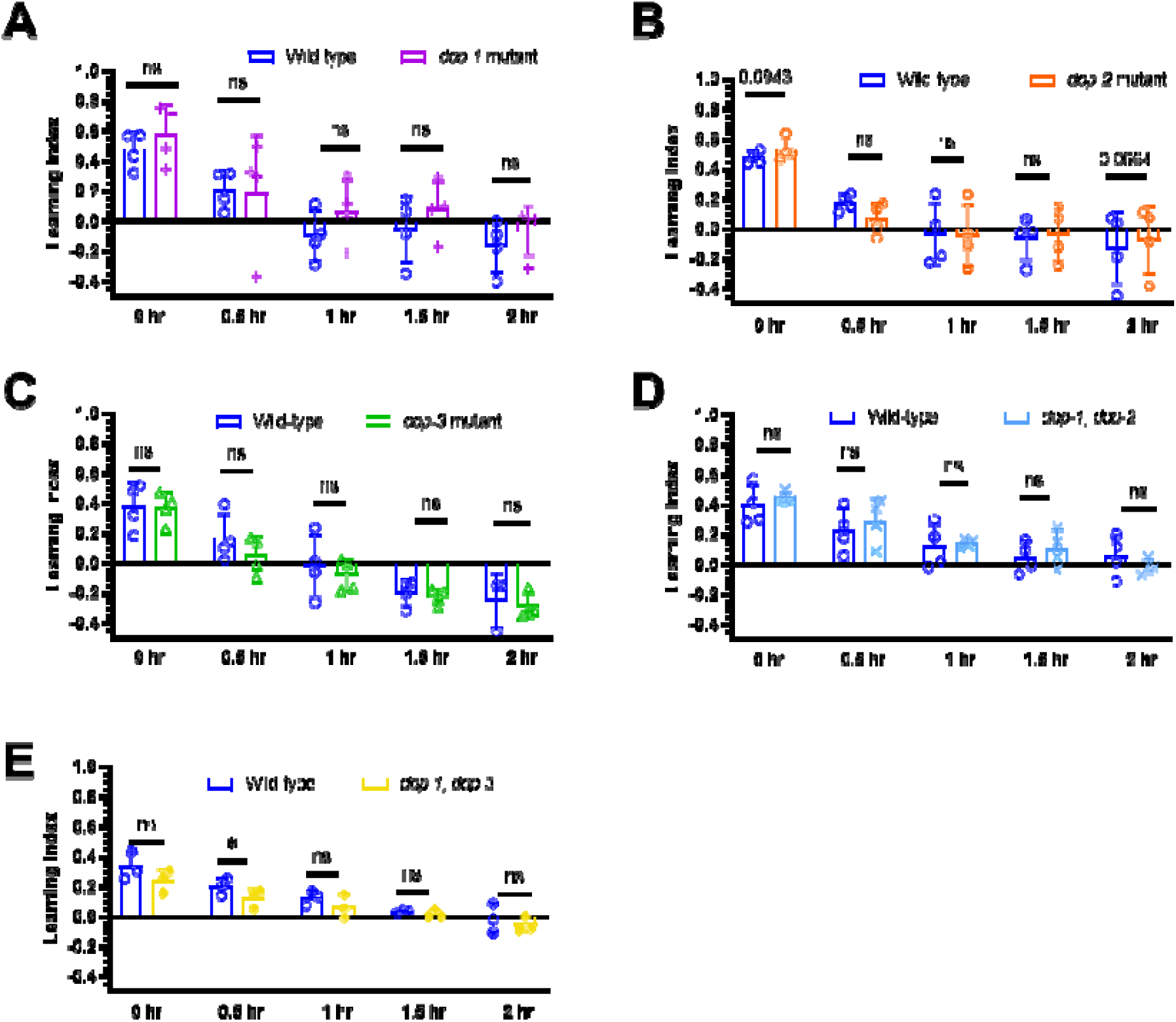
Dopamine receptors *dop-1*, *dop-2* and *dop-3* act redundantly to modulate short-term memory. Learning indices calculated for naïve/untrained worms and conditioned animals at 0, 0.5-, 1-, 1.5- and 2-hours post-conditioning, for chemotaxis data shown in Figure 2: **A)** *dop-1* mutants, **B)** *dop-2* mutants, **C)** *dop-3* mutants, **D)** *dop-1;dop-2* double mutants, **E)** *dop-1;dop-3* double mutants. Graphs show 3-4 biological replicates: each data point represents one biological replicate (which includes four technical replicates). Each technical replicate consists of 50-250 worms. Statistical analysis: a one-way ANOVA with Tukey’s multiple comparisons test was performed to compare between genotypes for all post-conditioning time points. Error bars = mean ± SD. p-values are represented by **** ≤ 0.0001; *** ≤ 0.001, ** ≤ 0.01; * ≤ 0.05; ns = non-significant.

