## Supplementary figures and images for "Dopaminergic Modulation of Short-Term Associative Memory in *Caenorhabditis elegans*"

### Supplemental Figure 1

#### Figure S1 (McMillen *et al.*)

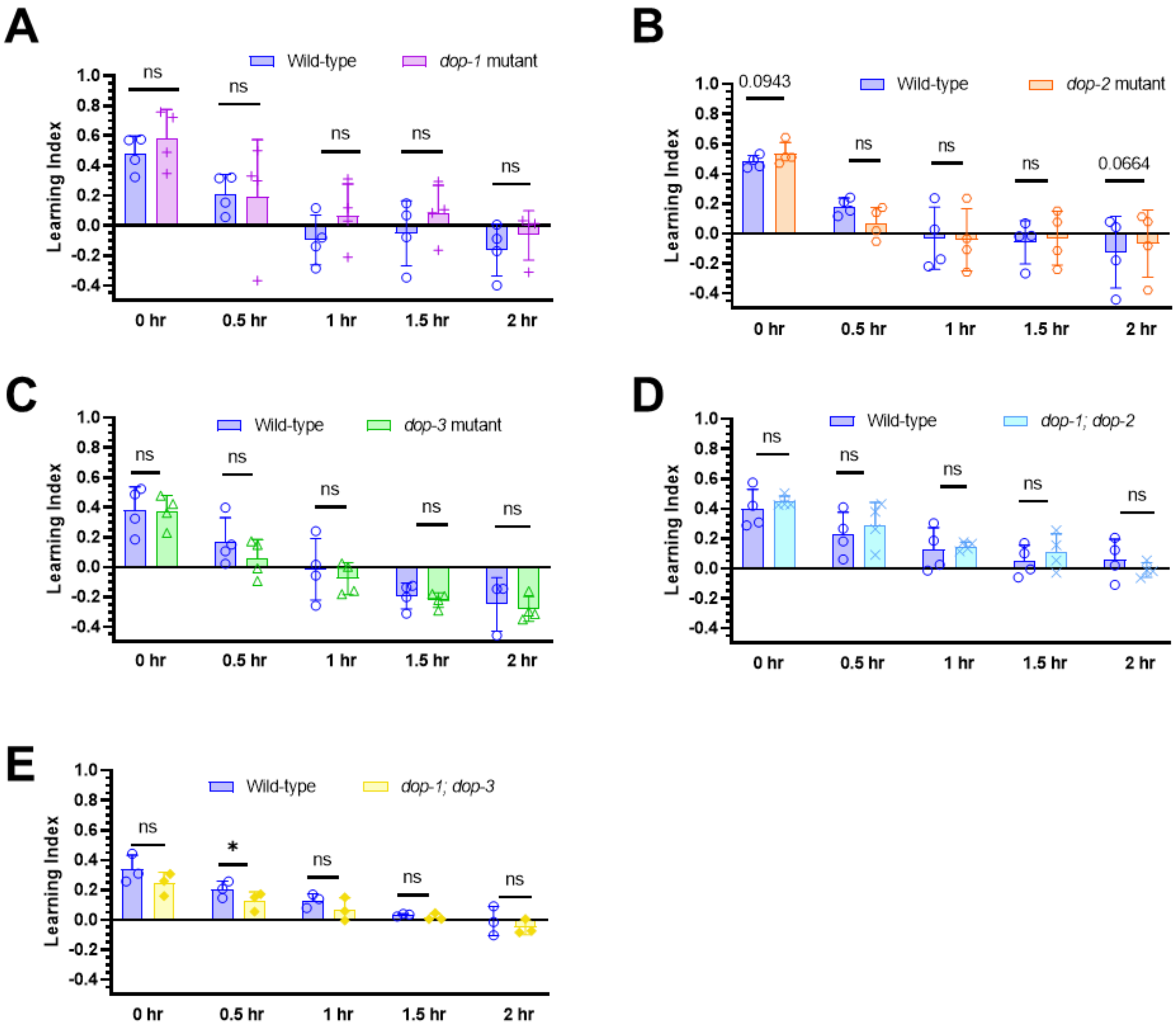
